# *Klebsiella pneumonia*, one of potential chief culprits of non-alcoholic fatty liver disease: through generation of endogenous ethanol

**DOI:** 10.1101/446765

**Authors:** Xiao Wei, Xiangna Zhao, Chen Chen, Jing Lu, Weiwei Cheng, Boxin Li, Huan Li, Weishi Lin, Changyu Tian, Jiangtao Zhao, Daizhi An, Juqiang Han, Xuejun Ma, Wei Li, Xuesong Wang, Xiao Chen, Zheng Zhang, Hui Zeng, Ying Sun, Ruifu Yang, Di Liu, Jing Yuan

## Abstract

Non-alcoholic fatty liver disease (NAFLD), a prelude of cirrhosis and hepatocellular carcinoma, is the most common chronic liver disease worldwide. NAFLD has been considerated to be associated with the composition of gut microbiota. However, causal relationship between change of gut microbiome and NAFLD remains unclear. Here we show that *Klebsiella pneumoniae* was significantly associated with NAFLD through inducing generation of endogenous ethanol. A strain of high alcohol-producing *Klebsiella pneumoniae* (HiAlc *Kpn*) was initially isolated from fecal samples of patient with non-alcoholic steatohepatitis (NASH) accompanied with auto-brewery syndrome (ABS). Gavage of HiAlc *Kpn* was capable of inducing murine model of fatty liver disease (FLD) in which had typical pathological changes of hepatic steatosis and similar liver gene expression profiles to those of alcohol intake in mice. Data derived from germ-free mice by gnotobiotic gavage further demonstrated that the HiAlc *Kpn* is the major cause of the changes in FLD mice. Furthermore, using proteomic and metabolitic analysis, we found that HiAlc *Kpn* induced generation of endogenous alcohol through the 2,3-butanediol fermentation pathway. More interestingly, the blood alcohol concentration was elevated in FLD mice induced by HiAlc *Kpn* after glucose intake. Clinical analysis showed that HiAlc *Kpn* were observed in up to 60% of patients with NAFLD. Our results suggested that HiAlc *Kpn* make important contribution to NAFLD, possibly through generation of the endogenous alcohol. Thus, targeting these bacteria might provide a novel therapeutic for clinical treatment of NAFLD.

**In Brief:** Fatty liver disease induced by high alcohol-producing *Klebsiella pneumoniae*

**Competing Financial Interest Statement:** The authors declare no conflicts of interest.

## Introduction

Fatty liver disease (FLD) is a chronic reversible disorder of liver with the hepatic manifestations of metabolic syndromes, which globally affects 10% to 24% of populations in various countries and up to 75% in obese subjects^1–3^. According to the etiology and the behavior of alcohol intake, FLD can be mainly classified as the nonalcoholic FLD (NAFLD) and the alcoholic FLD (AFLD)^4^. Both forms of the disease normally are initiated with fat deposition in liver and followed with liver injury, including steatohepatitis, inflammation, fibrosis, cirrhosis and hepatocellular carcinoma^45^. The major cause of the AFLD is alcohol intake, while that of the NAFLD remains unclear. Increased evidence has shown that NAFLD is strongly associated with obesity, the metabolic/insulin resistance syndrome, dyslipidemia^6–8^ and alterations of gut microbiota^9^.

Alterations of constitutional microbiota, such as *Firmicutes, Bacteroidetes, Actinobacteria*, and *Proteobacteria*, might impair the basic *in vivo* functions including the immune system, the maintenance of nutrition, xenobiotics metabolism, development and proliferation of intestinal cells, and protection against aggressor microorganisms. Metagenomic analyses revealed that the metabolic diseases such as obesity^10–13^, the metabolic syndromes^14^, non-alcoholic steatohepatitis (NASH) and cirrhosis^9^ are the results of disorder of the composition of gut microbiota. Particularly, it has been shown that the enrichment of *Eubacterium rectale, E. rectale, Bacteroides vulgates and etc*. correlates with NAFLD, possibly through effecting of harmful metabolic mediators on the host^15^. For example, a recent study reported that the endogenous alcohol generated by gut microbiota may affect the progress of NAFLD^16^. Hence, it is of particular interest to identify which bacteria is the major culprit that causes the development of FLD and to illustrate molecular mechanisms involved in the pathogenesis^9 17, 18^. Our inspiration came from a case of patient with severe NASH, who accompanied with auto-brewery syndrome and had elevated blood alcohol concentration after eating alcohol-free high-carbohydrates. Surprisingly, we found that this was due to bacterial rather than that of fungi because antifungal treatments did not have effects on such syndrome. Here, we reported as the first case of bacterial ABS, who eventually recovered after antibiotic treatments. In this case, we isolated and identified some strains of *Klebsiella pneumonia* which were highly associated with endo-alcohol producing (HiAlc *Kpn*). Considering that the NAFLD might be induced by endo-alcohol, we attempted to connect these commensal HiAlc *Kpn* and the pathogenesis for hepatic damage. Through gastric gavage of the HiAlc *Kpn*, we established a murine model of FLD, which confirmed HiAlc *Kpn* is as an important causative agent of FLD via induction of endogenous alcohol. Using this model, we exposed the pathogenesis of NAFLD and determined the molecular mechanisms of HiAlc *Kpn-mediated* ethanol fermentation. Finally, we also observed that this HiAlc *Kpn* widely presents in other NAFLD patients. Our findings might provide benefits for clinical treatments and for potential noninvasive methods to detect NAFLD.

## Results

### HiAlc *Kpn* strongly correlated with NAFLD

The metagenomic analysis of 16S rDNA was performed in 14 consecutive fecal samples collected from a patient with NASH/ABS, during the pre-onset, onset, recovery, and post-treatment stages (Fig. S1a). Total 738,865 sequence reads were obtained and were assigned mainly to seven bacterial phyla (Fig. 1a). When mapping the metagenomic data onto the curves for blood alcohol concentration (BAC), it was noticed that the distribution of the phylum *Proteobacteria* strongly correlated with fluctuations of BAC (R=0.89) (Fig. 1a). Intriguingly, the abundance of *Klebsiella* of *Proteobacteria* reached 18.8% in the first day of the morbid state, nine hundred-fold higher than the healthy controls (~0.02%) (Fig. 1a and Fig. S1b). In contrast, yeast test was negative in these samples.

**Figure 1.**
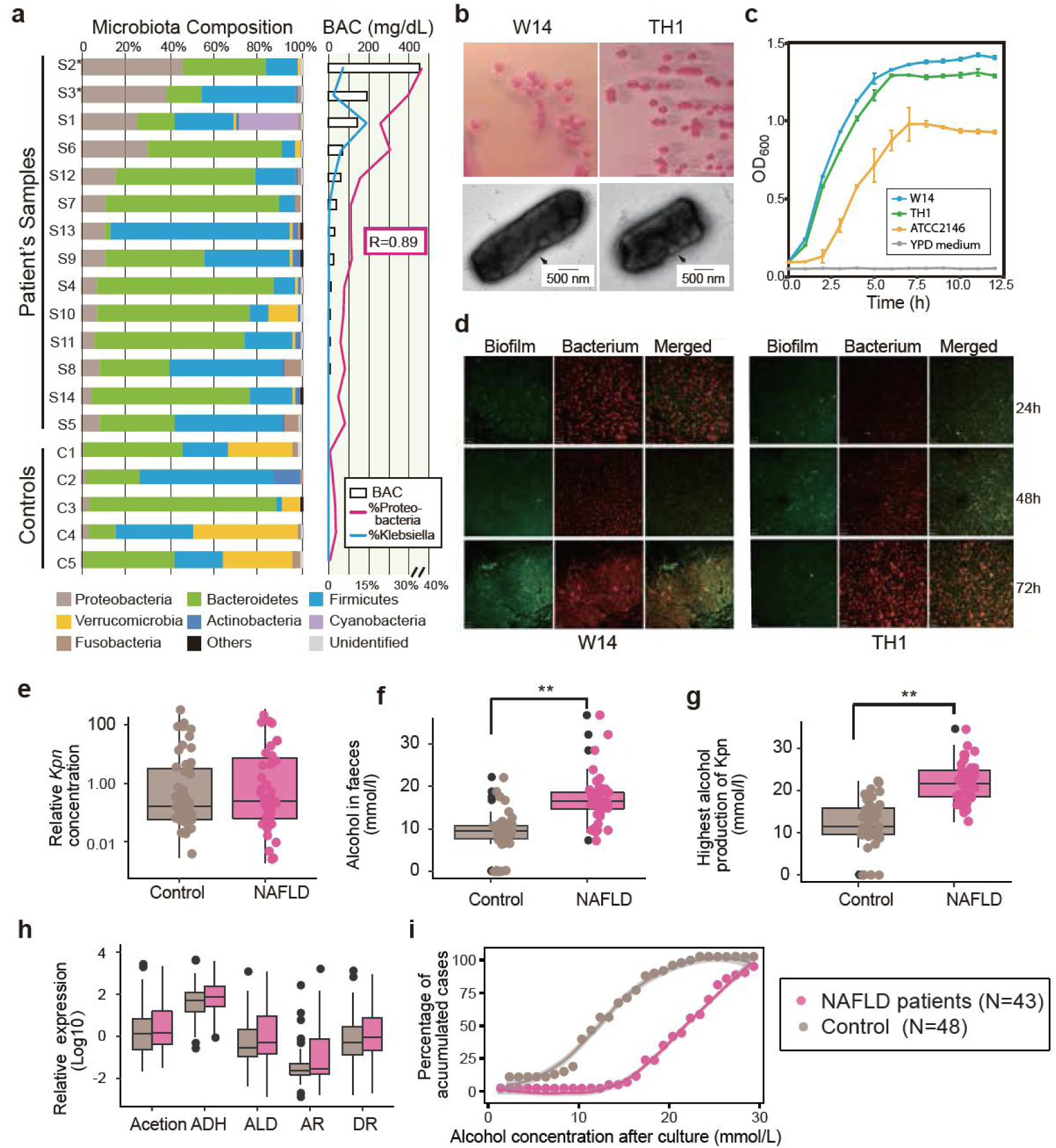
Commensal HiAlc *Kpn* has a higher statistical chance of initialing NAFLD. **(a)** Correlations of the NASH/ABS patient’s intestinal microbiota and blood alcohol concentration (BAC). Gut microbiota compositions of samples are placed according to the BAC (from higher to lower). The percentages of *Proteobacteria* and *Klebsiella*, respectively, are outlined. The correlations of *Proteobacteria* and BAC are calculated. Colonies and capsules images by TEM **(b)**, growth curves **(c)**, and Laser scanning confocal microscopy images to show the biofilm formation of *K. pneumoniae* W14 and TH1 isolated from the NASH/ABS patient. (**e-h**) The alcohol producing ability from *Kpn* in NAFLD population are higher than the controls. **e**, Relative *Kpn* quantification measured. (**f)** Alcohol producing ability was measured by detection the alcohol concentration in fermented fecal sample in NAFLD patients (brown box) and the controls (carmine box). (**g)** Alcohol producing ability was measured with the highest alcohol producing *Kpn* isolation. (**h)** Evaluated the alcohol producing genes. (i) Accumulated cases increased with *Kpn* alcohol producing ability, showing that the NAFLD patients has a higher chance containing HiAlic *Kpn*. Loess smoothing method was used to regress the relation between accumulated case and *Kpn* alcohol producing ability with grey shadows. Values are the mean ± SD obtained from multiple independent experiments. In panels: *P < 0.05, **P < 0.01 (unpaired *t*-test). In box plot, centerline indicates the median; box outlines show 25th and 75th percentiles, and whiskers indicate 1.5× the interquartile range. Extreme values are shown separately (black dot).

Using the yeast extract peptone dextrose (YPD) medium with 10% alcohol, we isolated two alcohol-tolerant strains of *K. pneumonia* (W14 and TH1) that produced the highest amounts of alcohol under both aerobic (63.2 and 60.8 mmol/L, respectively) and anaerobic conditions (36.7 and 31.2 mmol/L, respectively), which named HiAlc *Kpn*. These HiAlc *Kpn* strains appeared as typical mucoid lactose fermenters, having clear capsules and biofilms and higher growth speed (Fig. 1b-d). The *in vitro* cultivation experiments showed that ethanol production ability of these strains was related to both carbon source and air conditions (Fig. S1c).

To validate the correlation between HiAlc *Kpn* and NAFLD, we analyzed the abundance of *Kpn*, the ability of alcohol producing as well as the expression of genes associated pathways in patients with NAFLD (n = 43) and control subjects without FLD symptoms (n = 48). Results showed that the abundance and ability producing alcohol of *Kpn* were higher in the fecaes of NAFLD patients compared with those of healthy individuals (Fig. 1f-h). With increasing alcohol concentrations of culture medium, more alcohol tolerant bacterial clones with higher alcohol producing ability including *Kpn* were identified in patients with NAFLD compared with those of controls (Table S1 and Fig. 1h). In addition, key enzyme genes associated with alcohol producing pathways also expressed at higher levels in NAFLD patients compared with those of controls. In our cohort, 61% NAFLD patients carried HiAlc and MidAlc *Kpn* (the alcohol-producing concentration ≥20mmol/L), while that was only 6.25% in controls (Fig. 1i). This suggested that the HiAlc *Kpn* is highly associated with NAFLD.

### HiAlc *Kpn* induced murine model of FLD

To explore association of the HiAlc bacteria and the progression of FLD, we fed groups of the SPF mice with the HiAlc *Kpn* for 4, 6 and 8 weeks, while mice fed with ethanol and with YPD medium (pair-fed) were used as positive and negative controls, respectively (Fig. S2a). There were no significant differences in body weight and liver body mass ratio of mice in all groups studied (Fig. S2b and 2c). The histological staining showed that microsteatosis and macrosteatosis clearly presented in the livers of the HiAlc *Kpn*-fed mice at 4 and 8 weeks, respectively, which were comparable with those of changes in the ethanol-fed mice (Fig. 2a). However, histological and immunehistochemical stainings of Sirius Red and α-smooth muscle actin showed that there was no obvious liver fibrosis (data not shown). These suggested that the HiAlc *Kpn* feeding is capable of inducing development of hepatic steatosis in mice.

**Figure 2.**
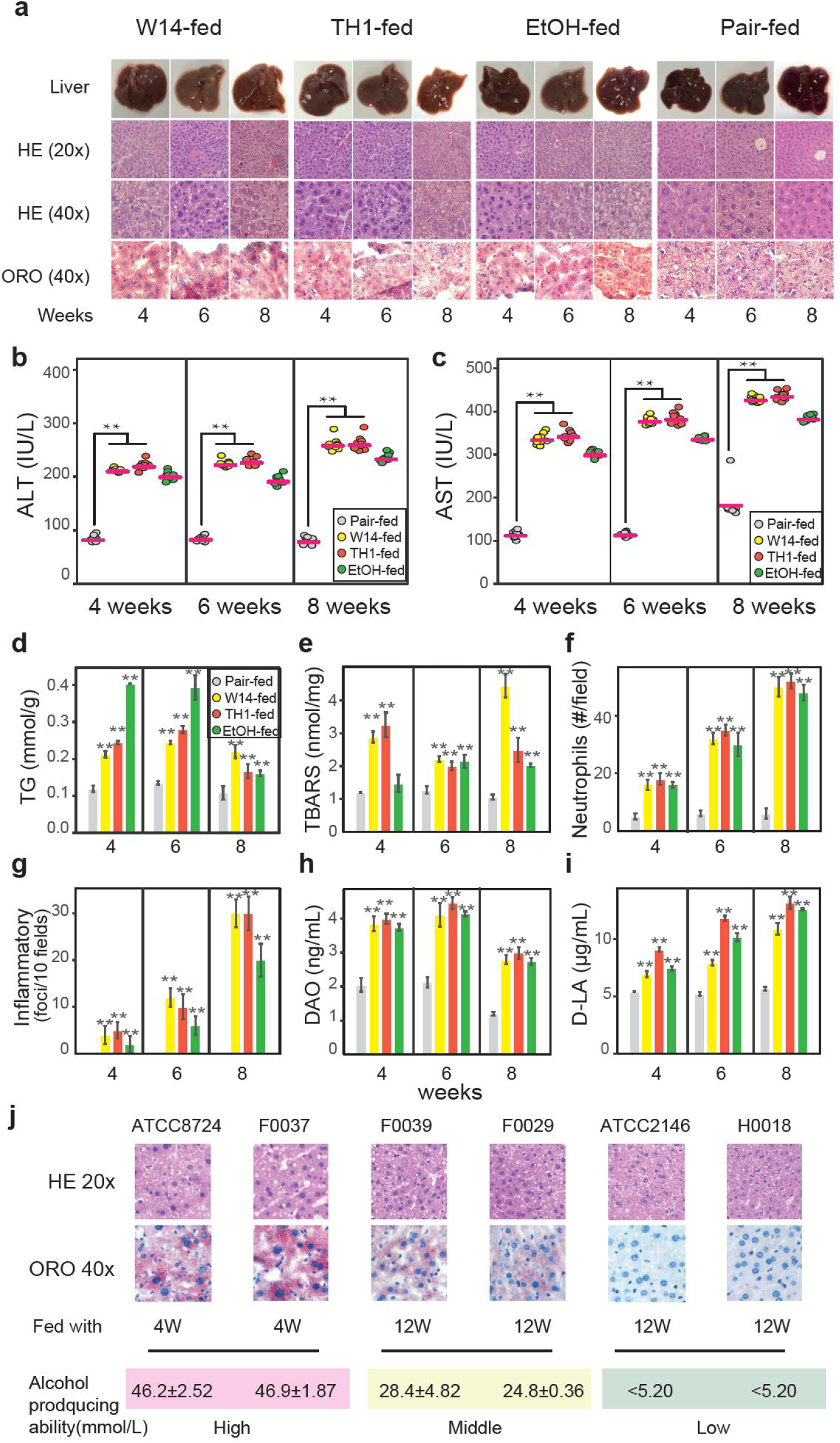
Mice feeding with HiAlc *Kpn* could establish a chronic hepatic steatosis model. **(a)** Anatomy, HE staining (20× and 40× magnifications), and Oil Red O staining (40× magnification) of SPF mice liver feeding with HiAlc *Kpn*, EtOH, and pair for 4, 6, and 8 weeks. (**b-i)** Liver injury and intestinal permeability of FLD mice induced by HiAlc *Kpn* feeding. The serum levels of ALT (**b**) and AST (**c**), and the contents of TG (**d**), TBARS (**e**), neutrophils (**f**), and inflammatory (**g**) in the liver tissues are used to assess liver injury. The serum levels of DAO (**h**) and D-LA (**i**) are used to assess the intestine permeability. (**j**) The level of alcohol producing ability resulted in different clinical outcomes. High (ATCC8724 and F0037), middle (F0029 and F0039) and low (ATCC2146 and H0018) alcohol producing isolates were fed to SPF mice. HE staining (20× magnifications), and Oil Red O staining (40× magnification) were used to evaluate the liver injure. SPF mice fed with HiAlc and middle alcohol producing *Kpn* occurs liver injure at 4 weeks and 12 weeks, respectively. However, the mice fed with low alcohol producing *Kpn* did not show live injure until 12 weeks. Values are the mean ± SD. In panels: *P < 0.05, **P < 0.01 (unpaired t-test). Values in this figure were obtained from multiple independent experiments.

Measurements of aspartate transaminase (AST) and alanine transaminase (ALT) in serum, and triglyceride levels (TG) and thiobarbituric acid-reactive substances (TBARS) in liver showed that the clinical indices significantly increased in HiAlc *Kpn* gavage groups compared with negative control mice (Fig. 2b-2e), which further indicated occurrence of the pathophysiological dynamic changes in livers of these mice. Besides, the numbers of neutrophils and inflammatory foci were also significantly elevated in HiAlc *Kpn-* and ethanol-fed groups with the time course of gavage (Fig. 2f and 2g).

Furthermore, HiAlc *Kpn-* and ethanol-fed FLD mice had remarkable morphological changes of intestinal villi (Fig. S2d), companying with significant increases of the levels of intestinal diamine oxidase (DAO, P<0.05) and D-lactate content (D-LA, P<0.05) compared to the pair-fed group (Fig. 2h and 2i), suggesting that the colonization of HiAlc *Kpn in vivo* might affect integrity of epithelium and intestinal permeability as alcohol did.

In addition, we compared effects of several other *Kpn* strains isolated from fecal sample (F0037, F0039, F0029 and H0018 with abilities of high, middle and low alcohol producing) with those of *K. oxytoca* ATCC8724 which were used in industry, in feeding mice. Same as what observed in W14/TH1, typical physiologic dynamic in mice liver and the hepatic steatosis occurred at 4 weeks in the HiAlc strains (Fig. 2j). In middle alcohol producing group, this damage did not occur until 12 weeks. However, these changes were not observed in low alcohol producing group (Fig. 2j).

### Confirmation of the link between HiAlc *Kpn* and FLD in germ-free mice

To assess whether the HiAlc *Kpn* is the agent caused in FLD, we used germ-free mice in our gavage model to exclude the potential impact from other symbiotic gut microbiota. After gavage for 4 weeks, we observed that such mice had a high level of the colonized HiAlc *Kpn* (Fig. 3). Histological staining with HE and Oil Red O showed clear hepatic steatosis, demonstrating that commensal HiAlc *Kpn* could initiate the FLD by producing endo-alcohol. Again, AST and ALT, TG, TBARS and the other clinical indices significantly increased in bacterial fed group (Fig. 3), which further supported that HiAlc *Kpn* is a key bacteria causing the progress of FLD.

**Figure 3.**
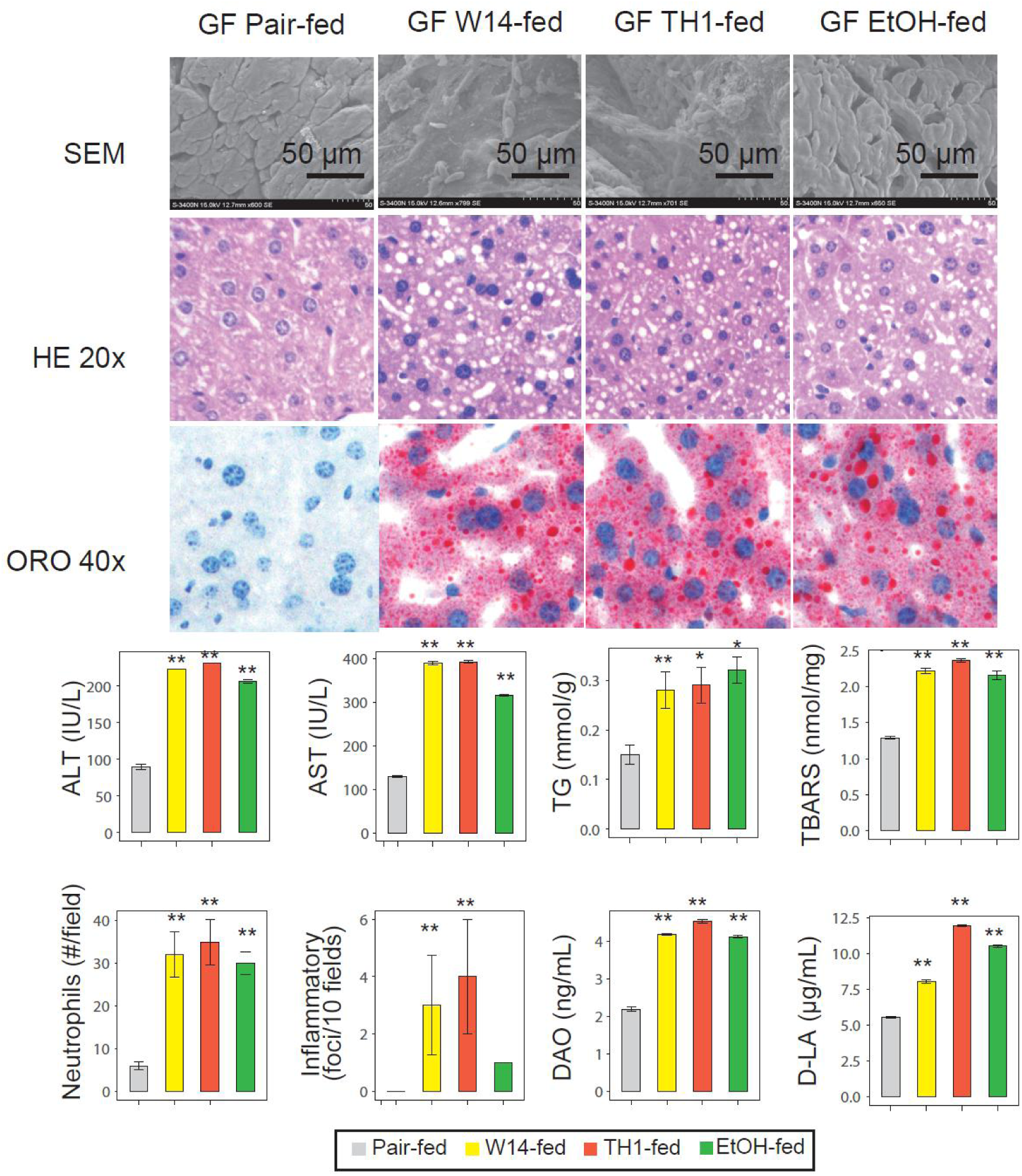
Confirming the link between HiAlc *Kpn* and NAFLD. Feeding with HiAlc *Kpn* was an important determinant for germ-free (GF) mice in FLD developments. Up panel, scanning electron micrograph (SEM) of the proximal colon, showing HiAlc *Kpn* the colonized status in vivo. Middle panel, liver histology and assessment of hepatic steatosis with HE staining (20× magnifications), and Oil Red O staining (40× magnification). Liver injury and intestinal permeability of GF mice measured by the levels of AST, ALT, TG, TBARS, neutrophils, inflammatory, DAO and D-LA. Values are the mean ± SD. In panels: *P × 0.05, ** < 0.01 (unpaired t-test). Values in this figure were obtained from multiple independent experiments.

### Alcohol producing pathway of HiAlc *Kpn*

To determine the molecular mechanisms and pathways of endogenous alcohol producing, we performed comparative proteomic and metabolic analysis in mice feces, *in vitro* and i*n vivo* cultivation in rabbits. Firstly, the results showed that 18 major metabolites were observed significantly increased in HiAlc *Kpn-* or ethanol-fed mice, including urea, alcohols, sugars, amino acids, and acids (Fig. S3a-3c; Table S2). Of special interest, 6 of these major metabolites were continuously elevated in HiAlc *Kpn-fed*, but none of them in ethanol-fed mice (Fig. S3a). Among these metabolites, the highest peak intensity was 2,3-butanediol, while citric acid and alpha-ketoglutaric acid were in higher concentrations in HiAlc *Kpn-fed* mice. Secondly, in vitro tests showed that the intensity of 10 metabolites, including 2,3-butanediol, ethanol and lactic acid were above 3.8e+006 (Fig. 4c). Among these, 2,3-butanediol and ethanol were in accordance with the abundant metabolites identified in the fecal samples of the HiAlc *Kpn-fed* mice. Thirdly, given that the difference of alcohol producing of HiAlc *Kpn* in aerobic and anaerobic conditions (Fig. S1c), we separated and identified the proteins that expressed in both conditions and with different abundance. Comparative proteomic analysis showed that 66 proteins were with 3-folds changes, consisting of 59 up-regulated and 7 down-regulated proteins (Table S3), while 21/59 proteins (account for 32%) were associated with the carbohydrate transport and metabolism pathway (Fig. 4a). In rabbit intestinal culture model, 10 of the 21 proteins mentioned above were identified, including the enzymes in the 2,3-butanediol fermentation pathway (Fig. 4b and Fig. S4a). These results suggested that the *in vitro* ability of HiAlc *Kpn* to produce alcohol might reflect the status of such bacteria *in vivo* (Fig. S4b). Finally, considering all potential alcohol producing pathways in bacteria, we found that a majority of the enriched proteins and metabolites were associated with 2,3-butanediol fermentation pathways (Fig. 4d), which normally is an ignored pathway in alcohol production from glucose and glycerol metabolism *in vivo*. Accordingly, the key up-regulated enzymes and metabolites were all occurred in this pathway (Fig. 4d).

**Figure 4.**
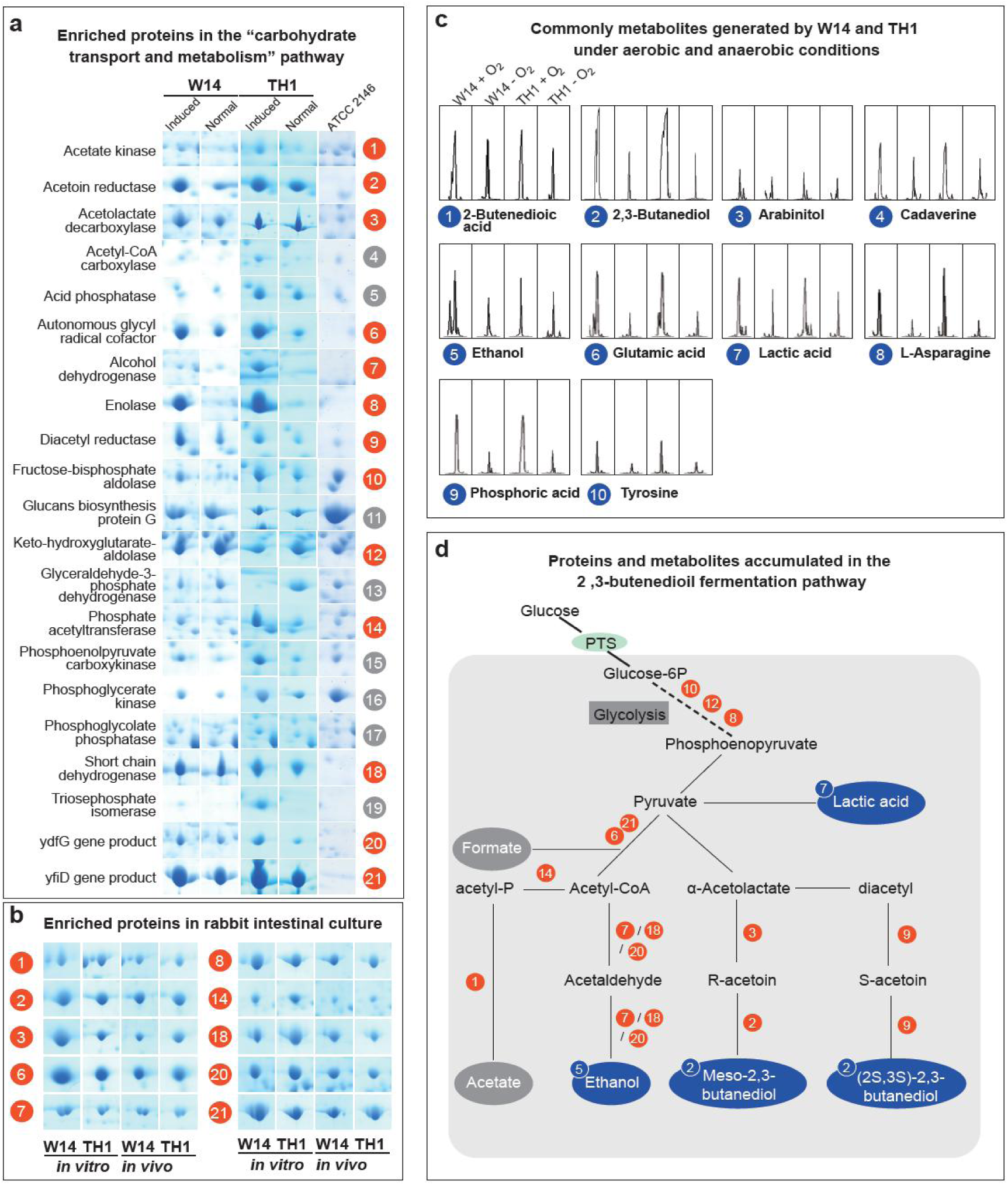
Proteomic and metabolomic analysis of HiAlc *Kpn* in aerobic and anaerobic conditions. **(a)**Significant proteins of HiAlc *Kpn* W14 and TH1 enriched in aerobic and anaerobic conditions. Twenty proteins up-regulated and one protein down-regulated are related to carbohydrate transport and metabolism, in especial the proteins covering 2,3-butanediol fermentation pathway. *Kpn* ATCC2146 is used as control. **(b)** Ten interesting proteins up-regulated of HiAlc *Kpn* W14 and TH1 grown to produce ethanol *in vivo* vs. *in vitro* are related to 2,3-butanediol fermentation pathway by using a model of a rabbit intestinal culture. **(c)** Common metabolites produced in HiAlc *Kpn* W14 and TH1 in aerobic and anaerobic conditions (Peak intensity ≥ 3.8e+006) further confirm the existence of the 2,3-butanediol fermentation pathway. **(d)** Proposed central metabolism of the 2,3-butanediol fermentation pathway in HiAlc *Kpn*. End products and intermediates are emboldened. The enzymes identified of the pathway are shown in red number and the metabolites identified by GC-MS are shown in blue cycles. The number with red or blue circles is according to the protein or metabolites identified in Fig. 4A and Fig. 4C.

### Liver gene expression dynamics in FLD progression

To further explore the overall development and progression of hepatic steatosis induced by HiAlc *Kpn*, we detected liver gene expression in HiAlc *Kpn-*, ethanol-, and pair-fed mice. After the gavage for 4 weeks, ten lipogenesis genes (*cyp4a10, adipoq, cyp4a14*, *srebp1c, scd1, acd, lxra, lxrα, fas, andlcad*) and one fat oxidation gene (peroxisome proliferator-activated receptor α, *PPARα*) were up-regulated in livers from the HiAlc *Kpn-* and the ethanol-fed mice than that in the pair-fed mice (Fig. 5 a). Following these, we examined liver transcriptional profiles during FLD progression at 4, 6, and 8 weeks respectively, using microarray technology. Compared with the negative control group, a large number of genes which expressed over 2-fold were mainly observed in the samples of the HiAlc *Kpn-* and ethanol-fed mice, after 4-week gavage, while samples of mice with 6- and 8-week gavage showed less differentially expressed genes (Fig. S5a-5b).

**Figure 5.**
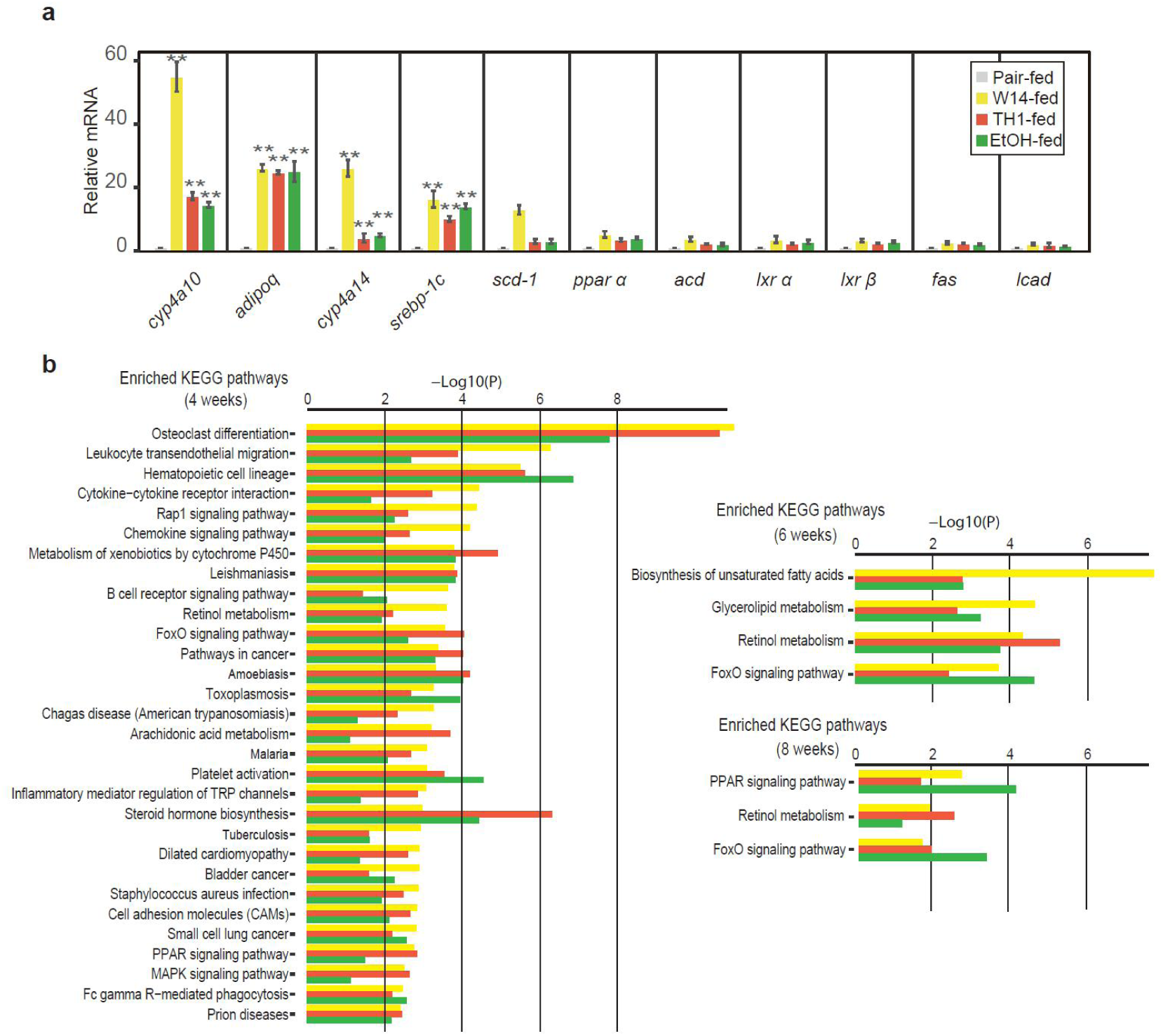
The key genes and biological processes during HiAlc *Kpn* induced FLD development. **(a)** Expressions of related genes to fatty-acid metabolism by using real-time PCR in mice liver after feeding by HiAlc *Kpn* for 4 weeks. **(b)** Enriched KEGG pathway of HiAlc *Kpn-* and EtOH-fed. Yellow, red and green are used to show HiAlc *Kpn* W14-, TH1-, and EtOH-fed, respectively. The negative 10-based logarithm of P-value is used to assess the significance.

Based on the differentially expressed genes, we further analyzed the enrichment on KEGG pathways, which occurred in both HiAlc *Kpn-* and ethanol-fed mice (Table S4). In the early stage (4 weeks), the common KEGG pathways were mainly involved, including osteoclast differentiation, retinol metabolism and arachidonic acid metabolism etc., which illustrated the progression of fat stored in liver cell with damage (Fig. 5b). In contrast, enrichments on biosynthesis of unsaturated fatty acids, glycerolipid metabolism, retinol metabolism, FoxO signaling pathway and PPAR signaling pathway might contribute to the process of the development to hepatic steatosis (6 and 8 weeks) (Fig. 5b). Notably, the pathways of retinol metabolism and FoxO signaling pathway were enriched in all three-time points observed, while enrichment of PPAR signaling pathway was observed in 4 and 8 weeks. These activated pathways have been proved to directly drive the increase of free fat acid (FFA), and cause FLD^19–21^. Furthermore, we also compared the enriched pathways between HiAlc *Kpn*-fed mice and the ethanol-fed mice, and found that a series of cancer-related pathways, including central carbon metabolism in cancer, colorectal cancer and p53 signaling pathway, were appeared after 4-week gavage, while only the p53 signaling pathway was appeared in the 8-week of the ethanol-fed mice (Fig. S6a).

In addition, analysis of Gene Ontology enrichment on the biological processes with the up-regulated genes highlighted the common biological processes for HiAlc *Kpn-* and ethanol-fed mice during FLD progress (Fig. S6b-6c). Apart from hepatic steatosis, the biological processes specifically enriched in inflammatory in the HiAlc *Kpn*-fed mice.

### Potential clinic diagnostic markers for HiAlc bacteria induced FLD

Based on our findings as above, we then attempted to discover the approach for endogenous alcohol detection and the potential marker for HiAlc bacteria induced FLD. We firstly compared the gut microbiota of HiAlc *Kpn*- or ethanol-fed mice against pair-fed mice, but did not find any accumulated bacteria populations that were related to alcohol absorption (Fig. S7). Then, considering the glucose is the substrate of alcohol producing, we tried to detect the blood alcohol concentration after fed with alcohol. i) The blood alcohol concentration was detectable in the ethanol-fed mice but not in those of HiAlc *Kpn-fed* mice without glucose gavage (Fig. 6). ii), However, the HiAlc *Kpn-fed* mice with glucose gavage had high blood alcohol concentration 2 hours after glucose gavage, which reached the peak value (111.16 mg/L in W14-fed mice and 75.65 mg/L in TH1-fed mice) at 4 hours (Fig.6), and the mice became inebriated. These implicated that the blood alcohol concentration following oral glucose intake might be considered as a clinical non-invasive diagnostic index for the detection of gut HiAlc bacteria, which might, therefore, further help the classification of FLD.

**Figure 6.**
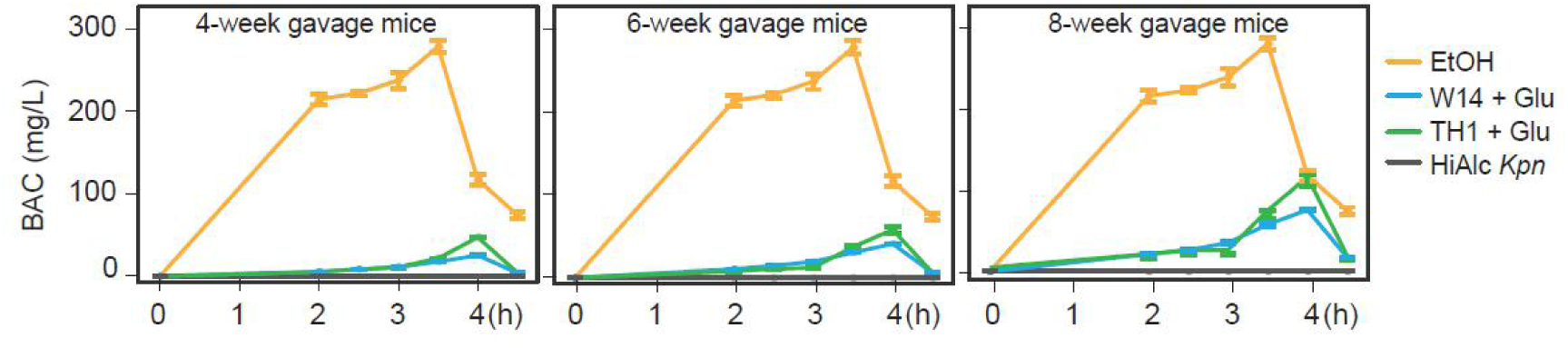
The blood alcohol concentration in HiAlc bacteria induced FLD mice. Dynamic changes of blood alcohol concentration of HiAlc *Kpn-fed* mice for 4, 6, and 8 weeks, and measured at 2, 2.5, 3, 3.5, 4 and 4.5 h after glucose inducing.

## Discussion

The liver-gut microbiota axis plays important roles in nutrition absorption and hepatotoxicity, within which the liver represents the first filter of nutrients, toxins and bacterial metabolites of blood supply from the intestine^22^. By aids of high-throughput techniques and improved bioinformatic tools, researches have been conducted to study the correlation of gene expressions in gut microbiota and host in the progress of NAFLD, and have proposed that gut microbiota was one of the important environmental factors affecting host metabolism and was highly associated with NAFLD^8,16^. Considering the variability between NAFLD patients and different metabolic pathways involved, however, how the gut microbiota cause hepaticsteatosis remains unclear. Herein, we established a murine model showing that the gavage of HiAlc *Kpn* strains, isolated from a rare NASH/ABS patient, directly induced FLD. These mice showed similar anatomic and histological appearances to the FLD mice-induced by alcohol-feeding, suggesting the endogenous alcohol production by gut bacteria could result in hepatic pathogenesis as those of alcohol intake. Our surveillance analysis of subjects with FLD indicated that a large proportion of NAFLD patients had endo-alcohol possibly because of HiAlc *Kpn*.

The “two-hit” hypothesis has been proposed to explain the pathogenesis of NAFLD/NASH progression^23^. The first hit, hepatic steatosis, is closely associated with lipotoxicity-induced mitochondrial abnormalities, while the second hit includes enhanced lipid peroxidation and increased generation of reactive oxygen species (ROS)^23^. Through the investigation of liver transcriptome, we identified enriched pathways of steroid hormone biosynthesis, biosynthesis of unsaturated fatty acids, PPAR, retinol metabolism, arachidonic acide metabolism that would increase the production of FFA, cause the dysfunction of mitochondria, and promote ROS in the HiAlc *Kpn*-induced mice. Moreover, a set of pathways of steatosis and inflammation were also unraveled to reflect the liver injury by constantly alcohol and ROS production. Those pathways discovered coincide with those found in the study of NAFLD^24^, implying the accordance with the “two-hit” hypothesis. Meanwhile, we also noticed up-regulated genes and enriched pathways related to alcohol catabolism, suggesting that excessive alcohol was scavenged from liver to counteract alcohol accumulation, and this accords with findings of AFLD progression^6^-^25^. The increased gut permeability of the HiAlc *Kpn-induced* mice also resembles the fact of AFLD mice. During the FLD progression, both HiAlc *Kpn-* and alcohol-induced mice showed the mechanisms of fatty infiltration in the liver and chronic inflammatory responses, and similar findings have previously been reported in both AFLD and NAFLD studies^5^.

These results pointed out that endogenous alcohol is very likely another risk factor for FLD. It is puzzling that patients with NAFLD and AFLD share many histologic features such as fat deposition around microvesicular and macrovesicular vessels and the indistinguishable number and size of Mallory bodies^4^. These similarities in the hepatic responses of NAFLD and alcohol exposure suggest that such conditions might evoke common pathogenic mechanisms. It is, however, conflicting whether the endogenous alcohol is the causative agent of NAFLD^26^. Some researchers have proposed that the elevation of endogenous alcohol is the result of insulin-dependent ADH impairment^27^, while others hypothesized that the endogenous alcohol is the culprit^5^. In the present study, using the HiAlc *Kpn* isolated from a patient’s gut, we clearly demonstrated that the endogenous alcohol produced by HiAlc *Kpn* could resulted in FLD in normal mice, while middle or low-level alcohol producing strains were hard to do so. Given observed other species in gut, such as *E. coli*, only having limit ability in alcohol production, which is hard to determine the roles in FLD progress and to observe the potential pathways impact by endo-alcohol *in vivo*. More important, HiAlc *Kpn* is not only occurring in a rare case, but widely in populations with FLD. Therefore, this FLD induced by HiAlc *Kpn*, through generation of endogenous alcohol, is different from AFLD and NAFLD reported previously, or at least, is a new type of NAFLD, namely Endo-AFLD.

Another mystery is that the HiAlc *Kpn* employ the 2,3-butanediol fermentation pathway to produce high-level endogenous alcohol, and this is different from the alcohol-production pathway used by yeasts. On the one hand, the pathway is capable of efficiently transforming sugar and glycerol into alcohols and acids. On the other hand, the metabolites during alcohol catabolism, including acetaldehyde, acetic acid and fatty acid ethyl esters, might also cause tissue injury and hepatic steatosis. However, the effects of other metabolites produced during 2,3-butanediol fermentation and catabolism are still unknown. Thus, the pathways to generate endogenous alcohol within intestine are to be investigated and the balance of endogenous alcohol production and conversion *in vivo* also requires to be clarified. Although it is widely accepted that low-carbohydrate, high-fat diet can lead to extreme weight gain and health risks such as obesity, single hepatic steatosis, and NASH^28^, our results added that the high-sugar diet might also increase the risk of FLD.

Given the high prevalence and increasing incidence of NAFLD, developments of early diagnosis approach are really required. Generally, alcohol is produced constantly by the intestinal microbiota in the human gut^29,30^. However, negligible blood alcohol concentration and the inability of gut microbiota to produce hepatotoxic concentrations of endogenous alcohol are undetectable in clinical diagnosis of obese, NAFLD, or NASH patients^9^. In the present study, we induced the higher blood alcohol concentration through feeding mice with high glucose- or fructose-containing food, which were also inebriated. Therefore, the use of oral glucose tolerance test (OGTT) might provide benefits for diagnosing Endo-AFLD patients and/or possibly ABS patients. Nevertheless, we did not observe any significantly increased gut bacteria populations that correlate with Endo-AFLD progression, although the genus *intestinimonas* significantly decreased. Clearly, additional comprehensive study needs to be done to further determine whether gut microbiota would be diagnostic markers.

Cause-and-effect relationships between gut microbiota and diseases have been recognized for obesity, inflammatory bowel disease, colorectal cancer, and etc.^12,13,31,32^. Data of the present study further support the modified Koch’s postulates^33^, and prove that HiAlc *Kpn* could result in Endo-AFLD. Moreover, the established mice model system would be further used for study of FLD and ABS. For the first time, we discovered the use of 2,3-butanediol fermentation pathway within intestine and determined the molecular mechanisms of alcohol production. More intriguingly, our study raised the potential connections of cryptic ABS and Endo-AFLD.

## Methods

### Correlation of HiAlic *Kpn* and BAC in FLD patients

A rare case of NASH accompanied with ABS (also known as “gut fermentation syndrome”) in a 27-year-old Chinese male, who became inebriated with ethanol concentrations in blood ≥190 mg/dL, even to ~ 400 mg/dL (the legal limit for alcohol in China is 20 mg/dL) using a Department of Transportation-approved alcohol breathalyzer after eating carbohydrates, he was even believed to be a “closet drinker”. In literature published previously, the underlying cause of ABS is thought to be an overgrowth of yeast that ferment carbohydrates into ethanol in the gut^34–37^. However, the patient was an anomaly because periodic auto-brewing was observed each month, and his illness did not subside after treatment with antifungal agents. Stool cultures and ITS rDNA PCR detection for fungi were also conducted, and the results were negative for yeast. The dynamic changes in intestinal microbiota community and the fluctuations in blood alcohol concentration were monitored from this patient. The activities of alcohol dehydrogenase (ADH) and aldehyde dehydrogenase (ALDH) in his body were 6.552 and 2.116 milli-units/mL, respectively, within normal ranges. The patient was then presented to a gastroenterology practice, where he underwent a complete gastroenterology workup. All results were negative. The patient denied having taken any types of yeast (such as probiotics) as nutritional supplementation and denied having any past history of gastrointestinal disorders or treatments. An EGD (esophagogastroduodenoscopy) and colonoscopy were conducted and the results were negative. The patient was then subjected to percutaneous liver biopsy, in which he fulfilled Kleiner’s criteria on hepatic fat infiltration, inflammation, fibrosis and Computed Tomography (CT) scan and the liver ultrasound, indicating NASH. Fecal samples from all specimens at different stages were subjected to metagenome DNA extraction, 16S rRNA sequencing, and metagenome sequencing, 5 healthy individuals were enrolled as control group.

### Isolation, identification and Biological characteristics of HiAlc *Kpn* in NASH/ABS patient

The high alcohol-producing strains were isolated using YPD medium with 5% alcohol under aerobic or anaerobic culture with the fecal samples during the morbid stage of the NASH/ABS patient. As for all isolates, PCR amplification and sequencing of 16SrDNA, microbial morphological analysis, electron microscope, MALDI-TOF mass spectrometry and automated microdilution techniques were used to identify bacterial colonies.

The growth curves of HiAlc *Kpn* W14 and TH1 were determined as previously described^38^. Alcohol-producing abilities of HiAlc *Kpn* were performed with YPD medium containing 2%, 4%, 6%, 8%, 10% fructose or glucose as the sole carbon source under aerobic or anaerobic condition, respectively. Standard strain *Kpn* ATCC2146 was used as control. Capsules of W14 and TH1 were observed by lactose fermenters on MacConkey agar and using Transmission electron microscope. Moreover, HiAlc *Kpn* were diluted 1:50 in YPD and cultivated in 6-well plates for 24h, 48h, and 72h, and biofilm forming was analyzed by Confocal Microscope.

### High-throughput sequencing of fecal microbiota and data processing

Fresh fecal samples (S1-S14) from different ABS stages, controls (C1-C5), and all mice (W14-, TH1-, EtOH-, and pair-fed) were collected and sequenced by using 16S rDNA gene V3-V4 region. In addition, fecal samples from patient at S3 and S9 stages were further analyzed by whole genome sequencing. Procedures for library generation, sequencing, and processing of longitudinal samples were as previously described^39^. Briefly, DNA samples were extracted using the QIAamp DNA stool Mini kit (Qiagen) following the manufacturer’s instructions. DNA library preparation was performed according to the manufacturer’s instruction (Illumina). Workflows were used to perform cluster generation, template hybridization, isothermal amplification, linearization, blocking and denaturation, and hybridization of the sequencing primers. Samples were run on an Illumina MiSeq for 2×250-bp paired-end sequencing^40^. The base-calling pipeline (version Illumina Pipeline-0.3) was used to process the raw fluorescence images and call sequences^40^. High quality reads were extracted by Mothurand Usearch, and assigned to taxonomy with QIIME (v 1.8.0). Whole metagenome data was assembled with the massively parallel short read assembler SOAPdenovo 2.20^41^, followed by performing the gene prediction by GeneMark v2.7^42^. All predicted genes were aligned pairwise using BLASTn. Genes, of which over 90% of their length can be aligned to another one with more than 95% identity (no gaps allowed), were removed as redundancies to construct a non-redundant gene catalogue.

### Bacterial Strains and Growth Conditions

Fecal samples of the patient closely for one incidence cycle were collected, cultivated and purified in both yeast extract peptone dextrose (YPD) and Maconkey medium(with or without 5% alcohol) under anaerobic and aerobic conditions at 37°C for 24h. Anaerobic condition was achieved in jars using AnaeroPacks (Mitsubishi Gas Chemical Company, Tokyo, Japan). Standard strain *K. pneumoniae* ATCC2146 was used as control.

### Measurement of alcohol concentration

Alcohol concentrations of all strains and fecael samples were measured with an ethanol assay kit from BioVision (Milpitas, CA), following the manufacturer’s instructions. The blood samples from the mice gavaged HiAlc *Kpn* were analyzed by headspace gas chromatography method (HS-GC, 6850 Agilent, with a flame ionization detector FID-Headspace)^30^.

### NAFLD patients

Moreover, we recruited fourty-three NAFLD patients and fourty-eight healthy volunteers who visited the Affiliated Hospital of Academy of Military Medical Science (AMMS) and Chinese PLA General Hospital in China for their annual physical examination. The liver imaging and liver biochemistry results of all NAFLD patients were hepatic steatosis whereas healthy controls were in the normal range. Physical examination, routine examination of blood, urine and stools, preoperative serological tests (including the detection of hepatitis B surface antigen, hepatitis C virus antibody, Treponema pallidum antibody, human immunodeficiency virus antibody), liver function, renal function, electrolyte, liver ultrasound, electrocardiogram and chest X-ray results were checked in all NAFLD and controls. Exclusion criteria included hypertension, diabetes, obesity, metabolic syndrome, IBD, alcoholic fatty liver disease, coeliac disease and cancer. The fecal samples from those groups were subjected to collect, followed by alcohol-producing ability detection and bacterial isolation. This study was approved by the Institutional Review Board of Affiliated Hospital of AMMS. All participants signed an informed consent form prior to entering the study. The study conformed to the ethical guidelines of the 1975 Declaration of Helsinki.

### Validation the correlation between HiAlc *Kpn* and NAFLD

Metagenomic DNA was extracted from fecal samples of NAFLD patients (n=43) and controls (n=48), and the abundance of specific gene *rcsA* (capsular polysaccharide synthesis regulating gene) of *Kpn*, and alcohol-metabolism related enzymes, including Acetion, ADH, ALD, AR, and DR were determined by RT-PCR analysis. To determine the relative expression of genes and avoid nonspecific reaction with the target gene from microorganisms, plasmid pGEX-BOT (36.2pg/μl) carrying with special oligonucleotide sequence from a botanic gene was used as internal standard (Ct = 30).

A total of 200 mg dry fecal samples from NAFLD and control objects were washed and fermented anaerobically in 100mL YPD medium for 12h, alcohol concentrations were detected in mid-exponential phase at an A_600_ of 0.9 corresponding to 1.5×10^8^ CFU/ml. The highest alcohol producing *Kpn* isolation of each case was selected and detected its alcohol production after cultured for 12h by using elevated alcohol tolerant experiment in Maconkey medium with 0%, 5%, and 10% alcohol, respectively.

### Construction of FLD mice model with strain gavage

Bal B/C Germ-free mice and C57BL/6J SPF mice were fed with a nutritionally adequate diet of standard laboratory chow for 5 days, then randomly divided into four groups and followed by gavaged once every two days at 10 mg/100 g body weight (~10^7^ CFU/mL *K. pneumoniae*, 300 μL) for 8weeks: HiAlc *Kpn*-fed groups were gavaged a single doses of *K. pneumoniae* W14 or TH1 suspended in YPD medium (~10^7^ CFU/mL, 300 μL), ethanol groups as positive control were gavaged a single doses of ethanol (40% ethanol, 300 μL), while pair-fed mice in negative control groups were gavaged YPD medium (300 μL). The gavage was always performed in the early morning. After gavage, mice were kept on control or ethanol diet and kept in the cages on the warm blanket with circulating water. 70% mice survived after strain or ethanol feeding. Following gavage, mice were slow-moving, but conscious and regained normal behavior within 4-6 h. The mice were always euthanized 9 h post gavage. Moreover, high *(K. oxytoca* ATCC8724 and F0037), middle (F0029 and F0039) and low *(Kpn* ATCC2146 and H0018) alcohol producing isolates or ATCC standrad strains were fed to C57BL/6J SPF mice according to the same method.

Continue to monitor the mice weight every weeks. After 4weeks, 6weeks, and 8weeks, the fresh feces of mice were taken for 16S rRNA and metabolomics analyses. Then, the mice were dissected to detect serum indies including ALT, AST, TG, TBARS, neutrophils, and inflammatory. The liver and small intestine were collected for pathological section. The serum levels of ALT and AST, and the contents of TG, TBARS, neutrophils, and inflammatory in the liver tissues are used to assess liver injury. The serum levels of DAO and D-LA are used to assess the intestine permeability.

### Histology and physiological assays in mice

Harvested Liver were fixed in 10% formalin and processed for H&E and Oil Red O staining. The serum was collected to measure blood chemistry (ALT, AST), liver TG and TBARS content, hepatic lipid contents, blood ethanol concentration, serum cytokine levels, real time PCR, lipid peroxidation, and intestine DAO and D-LA content of gut permeability.

To detect hepatic expression of several lipogenesis genes, specific primers targeting the interest genes were used by real-time qPCR. Briefly, quantitative PCR experiments were performed with a Light Cycler 2.0 PCR sequence detection system by using the Fast Start DNA Master SYBR Green kit (Roche Diagnostics). Melting-point-determination analysis allowed the confirmation of the specificity of the amplification products. The copies numbers of target genes from each sample was calculated by comparing the Ct values obtained from the standard curves with the Light Cycler 4.0 software. Standard curves were created by using a serial 10-fold dilution of DNA from pure cultures, corresponding to 10,^1^–10^10^ copies/g feces. The data presented are the mean values of duplicate real-time qPCR analyses.

### Mice and Rabbits

Male Germ-free mice of 10-12 weeks were obtained from a breeding colony at the animal facility of the Third Military Medical University. Eight- to ten-week-old male Specific-pathogen-free (SPF) C57BL/6J mice were maintained at Academy of Military Medical Sciences in accordance with Academy of Military Medical Sciences Animal Resource Center and the Institutional Animal Care and Use Committee (IACUC) guidelines. Six adult male Japanese white rabbits with ages ranging from 28 to 32 weeks were provided and kept by Experimental Animal Centre of AMMS. The mean body weight of the rabbits was 3,549 g (±203g).

### Metabolome Analysis by GC-MS

Fresh fecal samples from FLD mice induced by HiAlc *Kpn* and cultures of HiAlc *Kpn* were all subjected to metabolome analysis, which were preconditioned according to the manufacturer’s instructions. Briefly, 100 μL of each supernatant of fecal sample was spiked with an internal standard (10 μL ribitol solution, 0.2 mg/mL in H2O) and vortexed for 30s. The supernatant was dried under a stream of N2 gas. The residue was derivatized using a two-step procedure. First, 40 μL methoxyamine hydrochloride (20 mg/mL in pyridine) was added to the residue and shaken at 30°C for 90 min followed by 40 μL MSTFA (1% TMCS) incubated at 37 °C for 30 min. The samples were kept at room temperature for another 120 min, then stored at 4°C before injection.

GC-TOF/MS analysis was performed using a LECO Pegasus 4D system (Leco Corporation, St Joseph, MI), consisting of an Agilent 7890 gas chromatography coupled to a Pegasus 4D time-of-flight (TOF) mass spectrometer, with a DB-5 MS column (30 m×250μm i.d., 0.25μm, Agilent J&W Scientific, Folsom, CA, USA). The inlet temperature was 250°C. The carrier gas was helium kept at a constant flow rate of 1.0 ml/min. The GC temperature programming was set to 1 min isothermal heating at 70°C, followed by 5°C/min temperature ramp to 280°C, and held for 10 min. The transfer line and ion-source temperatures were 250°C and 220°C, respectively. Electron impact ionization (70 eV) was set at a detector voltage of 1,575 V. Ten scans per second were recorded over the full mass range of 50-800 m/z. Chromatogram acquisition, library research, and peak area calculation were performed using the ChromaTOF software (Version 4.5, LecoCorp.). Significantly different molecules were selected by FDR-adjusted *P* values.

### Proteomics analysis of HiAlc *Kpn* bacteria *in vivo* and *in vitro*

In vivo/in vitro culture assay by a rabbit intestinal model was performed as described previously^43^. HiAlc *Kpn* cells were diluted 1:50 in YPD and grown to an OD_600_ of 1.0 corresponding to 1.5 × 10^8^ colony forming units/mL. The bacterial culture was washed twice with prewarmed RPMI and resuspended in YPD medium. An amount of 20 mL of bacterial suspension was placed in dialysis tubing with a molecular weight cutoff of 20,000 Da (for interchange of the smaller host signal proteins/molecules in the intestine). After the rabbits were anesthetized, the HiAlc *Kpn* culture was implanted aseptically within the colon through a 1 cm incision, then the incision was closed using surgical staples. The tubing containing the HiAlc *Kpn* culture was either incubated in rabbit intestine for four hours or in vitro at 37°C. The rabbit generally was ambulatory within 4 h. Then, the dialysis bag containing HiAlc *Kpn* culture was took out and HiAlc *Kpn* was harvested for alcohol determination and proteomics analysis. The experiment was performed at least six times.

Whole cellular protein extracts was prepared. 2D gel electrophoresis or LC-MS/MS was carried out. Proteins were considered differentially expressed if their relative intensity differed more than 3-fold between the two conditions compared. Each experiment was performed at least three times. MALDI-TOF/TOF MS/MS measurements and Electrospray ionization MS/MS were performed to identify the proteins. MALDI-TOF/TOF MS/MS measurements were performed on a Bruker Ultraflex III TOF/TOF-MS (Bruker Daltonics GmbH, Bremen, Germany) equipped with a 337-nm wavelength nitrogen laser (model LSI 337i; Bruker) working in reflection mode. Electrospray ionization MS/MS was carried out with a hybrid quadrupole orthogonal acceleration tandem mass spectrometer (Q-TOF2; Micromass, Manchester, UK). MS/MS peak lists were created by MaxEnt3 (Mass Lynx v3.5; Micromass), and amino acid sequences were interpreted manually using MassSeq (Micromass). Peptide mass fingerprinting searches and all of the MS/MS ion data base searches were performed by using the program Mascot v2.2.06 (Matrix Science Ltd.) licensed in-house against the publically available Uniprot-Enterobacteriaceae database.

### Microarray-Based Gene Expression of HiAlc *Kpn-* fed Mice Liver

Whole genome expression analysis was performed by Shanghai OE Biotech Co., Ltd according to the protocol of one-color microarray-based gene expression analysis from Agilent Technology. The Agilent SurePrint G3 Mouse GE Microarray (8*60K,Design ID:028005) was used in this experiment. Total RNAs were prepared from livers of mice gavaged for 4weeks, 6weeks and 8weeksby RNeasy mini kit(QIAGEN). The sample labeling, microarray hybridization and washing were performed based on the manufacturer’s standard protocols. Genespring (version13.1, Agilent Technologies) were employed to finish the basic analysis with the raw data. Differentially expressed genes were then identified through fold change as well as *P* value calculated with t-test. The threshold set for up- and down-regulated genes was a fold change>= 2.0 and a *P* value<= 0.05. Afterwards, GO analysis and KEGG analysis were applied to determine the roles of these differentially expressed mRNAs.

### Time-cause analysis of sugar-riched diet for mice

In the next morning after fed by HiAlc *Kpn*, 300 μL of 10% glucose was gavaged to induce alcohol produce in HiAlc *Kpn-* fed mice, ethanol groups as positive control were gavaged a single doses of ethanol (40% ethanol, 300 μL), and pair-fed mice as negative control. The ethanol concentration in blood samples from the mice induced was measured at 2, 2.5, 3, 3.5, 4 and 4.5h by HS-GC.

### Statistical Analysis

ANOVA and Fisher’s tests were performed with R version 3.4.1. Data are expressed as means ± SD. *P* value less than 0.01 was considered statistically significant.

### Data Availability

The whole genome sequences of *K. pneumoniae* W14 and TH1 have been deposited at GenBank under accession number NZ_CP015753.1 for W14 and accession number NZ_CP016159.1 for TH1. The microarray information and data of mice liver sample are available at NCBI Gene Expression Omnibus (GEO) databases under the following accession: GSE102489. The raw illumina reads data of 16S rDNA and whole metagenome data for all samples from the patient and mice has been deposited in the NCBI Sequence Read Archive under accession number SRR5934751 and SRR5934662, respectively.

**Supplementary Information** included four tables and seven Supplementary Figures.

## Acknowledgements

We are grateful to Weijun Chen and Jun Yu for helpful advice; Hong Wei and Jie Liu of The Third Military Medical University for gnotobiotic mice assays; and YanPing Luo of Affiliated Hospital of Academy of Military Medical Science and Yiming Li of Chinese PLA General Hospital for providing the feceal samples from NAFLD patients and controls. This work was supported by a grant from the National Natural Science Foundation of China (31670035, 31370093 and 81790632) to J.Y., and Mega-projects of Science and Technology Research of China Grant 2017ZX100102.

## Author Contributions

J.Y., D.L., C.C., and Y.-R.F. led and conceived the project, designed and performed most experiments, analyzed and interpreted the data. J.L., W.-W.C., and B.-X.L. performed animal experiments and analyzed data. C.-Y.T., D.-Z.A., and X.-J.M. collected samples and performed clinical study. X.C. and H.Z. performed bacterial growth experiments. X.-S.W., Z.Z. performed DNA extraction experiments, 16S sequencing and data analysis. W.L., J.-Q.H., H.L., and W.-S.L. performed and analyzed proteomics and metabolite analysis. X.W. and X.-N.Z. performed microarray-based gene expression analyses. C.C., D.L. and J.Y. wrote the paper.

---

**Figure S1.**
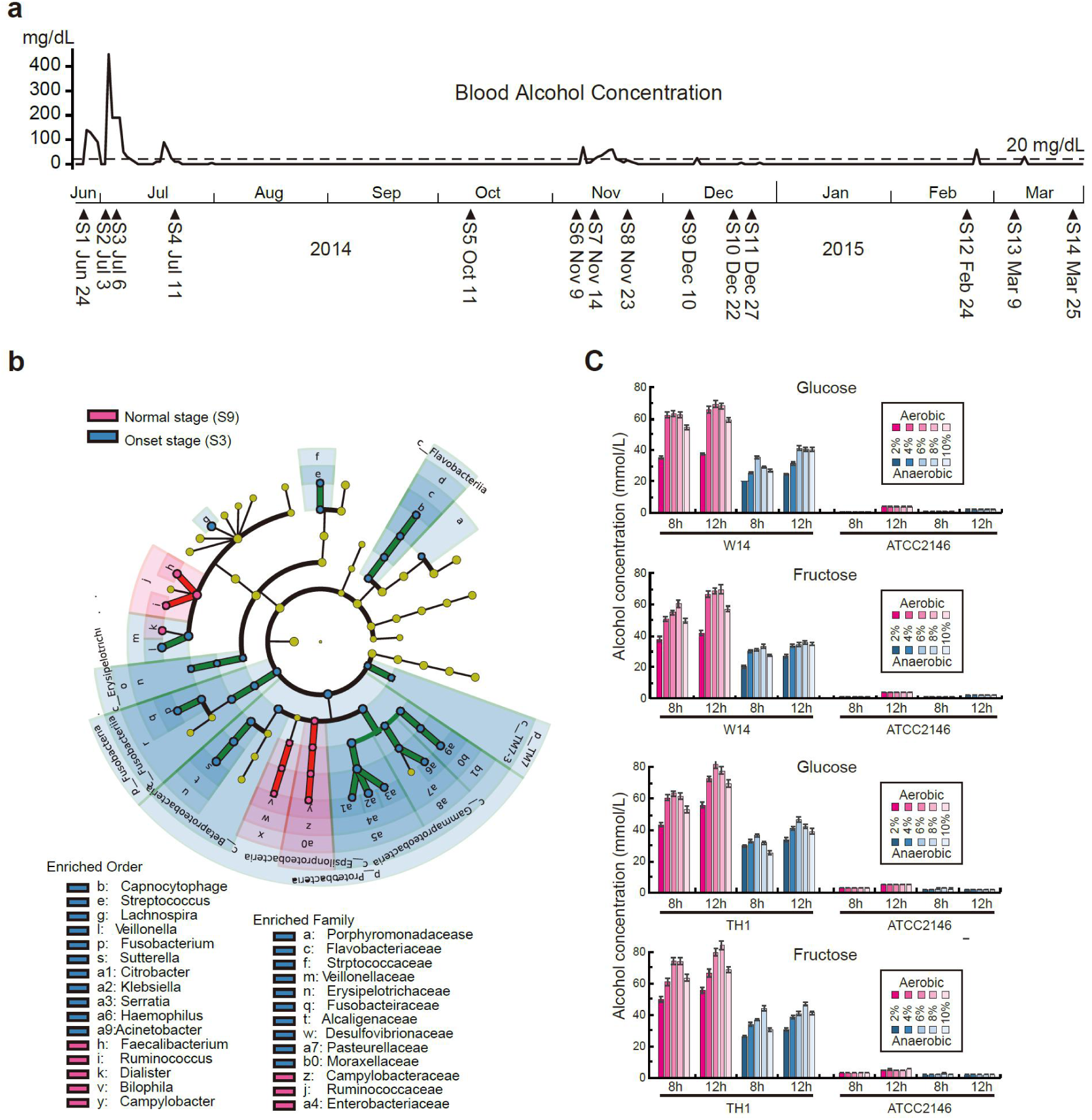
Timeline for the fecal samples of the patient collected and the whole genome LEfSe analysis of the NASH/ABS patient. **(a)** The times of the fecal samples from the patient collected in different stages and observation of blood alcohol concentration (BAC). The date was marked as from S1 to S14. **(b)** A cladogram representation of data in NASH/ABS patient in recovery stage and onset stage. Taxa enriched in onset (Green) and recovery control (Red). The brightness of each dot is proportional to its effect size. **(c)** Alcohol concentration of *Kpn* W14 and TH1 in YPD medium with sole carbon source (fructose or glucose) under aerobic and anaerobic conditions, respectively.

**Figure S2.**
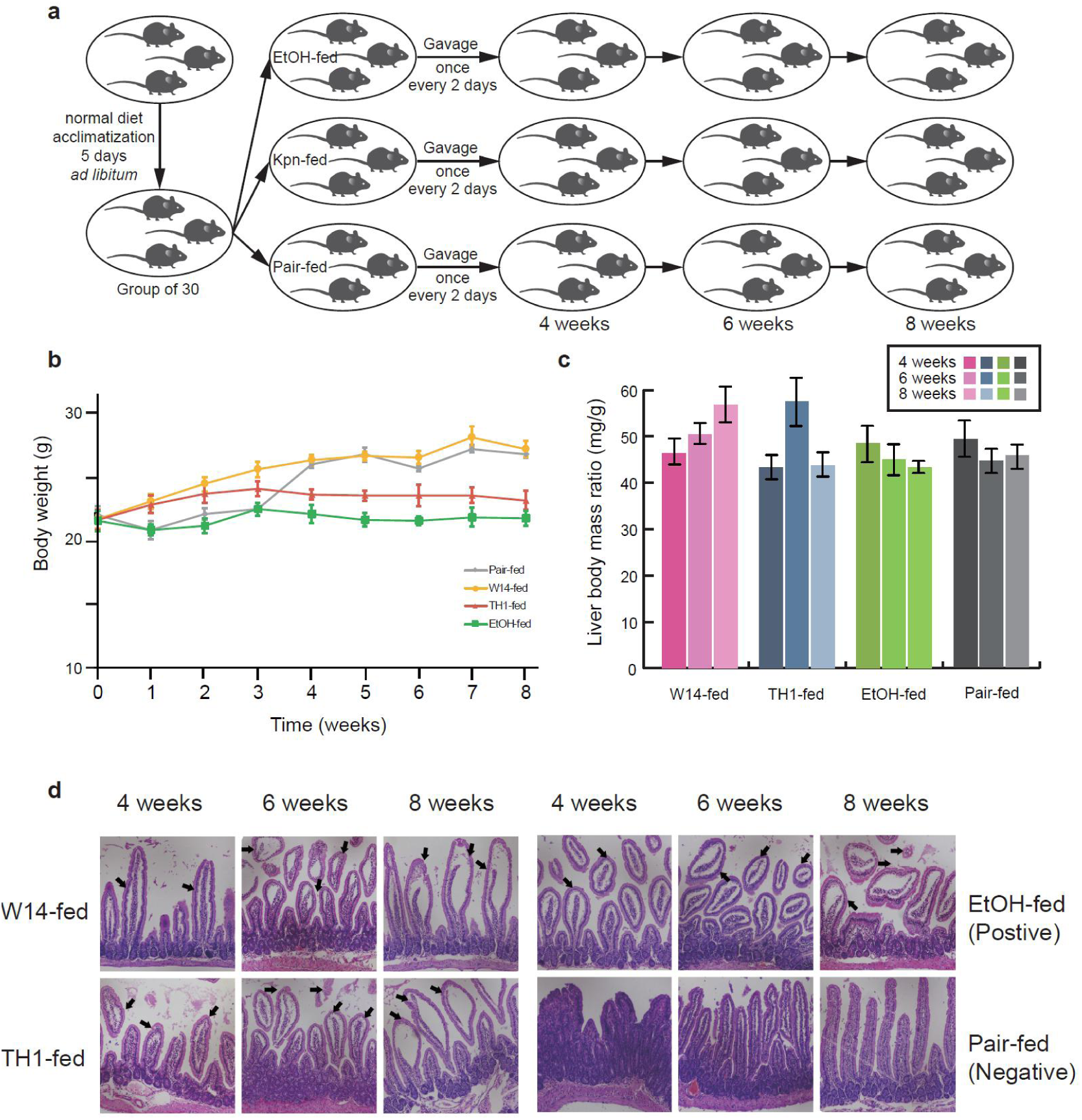
Physiological changes of FLD mice induced by HiAlc *Kpn*. **(a)** Schematic presentation of mice feeding. **(b)** Body weight changes of FLD mice during the HiAlc *Kpn* feeding. Data represent means ± SD. **(c)** Liver to body weight ratios of HiAlc *Kpn-*, EtOH-, and pair-fed mice at 4, 6, and 8 weeks. **(d)** Intestinal injury of FLD mice induced by HiAlc *Kpn* feeding. The histologic lesions of FLD mice intestine. The arrows in HE staining (20× magnifications) indicated the remarkable atrophy, edema, and shedding of intestinal villi in HiAlc *Kpn-* and EtOH-fed FLD mice, compared with pair-fed group (P<0.05).

**Figure S3.**
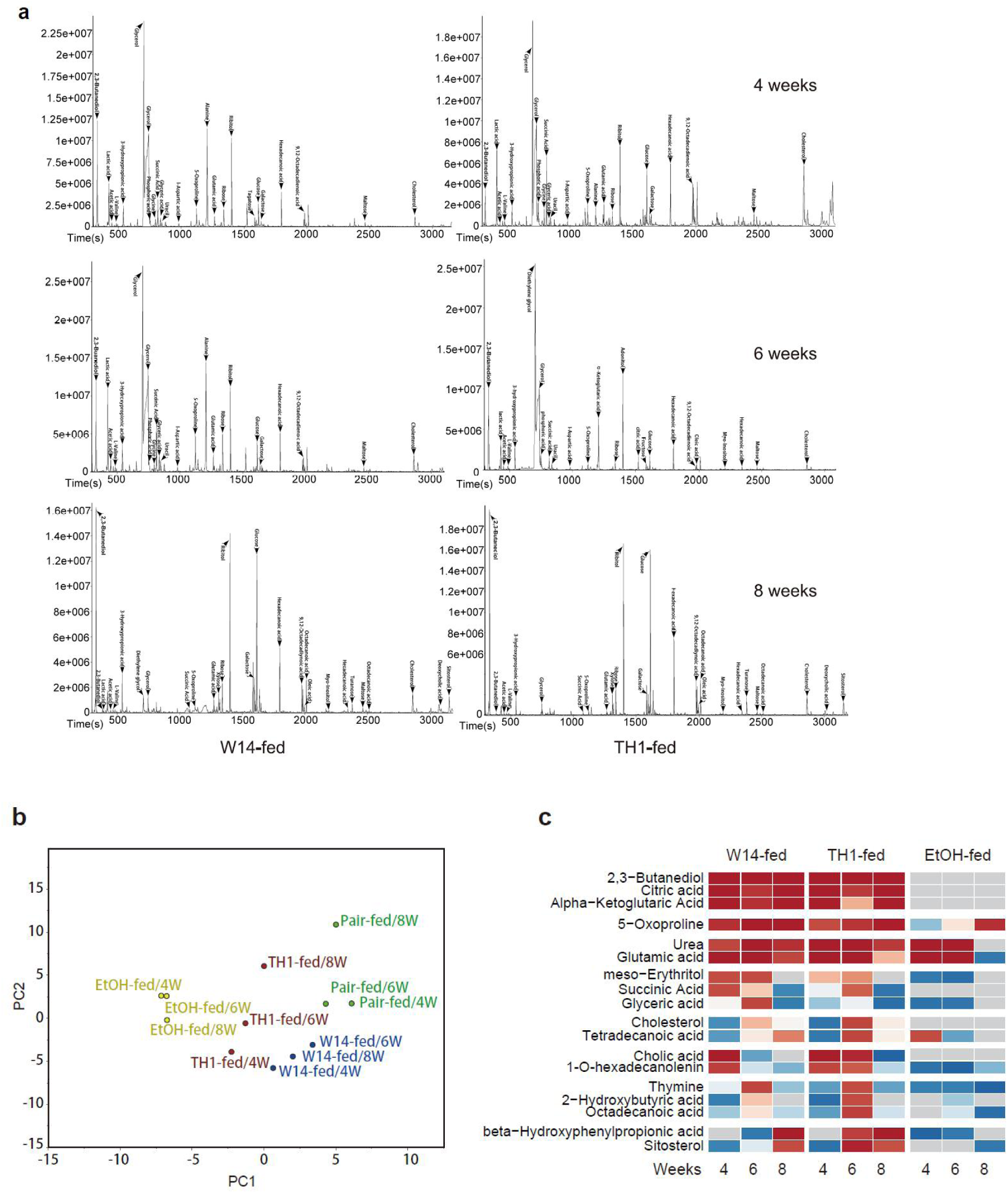
Metabolomic profilings of the fecal samples from FLD mice induced by HiAlc *Kpn* W14 and TH1 feeding, VOCs identified by GC-MS were marked in the peaks. **(a)** Metabolomic profilings of the fecal samples from FLD mice induced by HiAlc *Kpn* W14 and TH1 feeding. **(b)** PCA analysis of FLD mice induced by HiAlc *Kpn-*, ethonal-, and pair-fed at 4, 6, and 8 weeks. **(c)** Comparative metabolomic analysis of HiAlc *Kpn*-inducedYPD and grown to an FLD mice. Ratios of metabolite intensity in HiAlc *Kpn-* and EtOH-fed mice (against pair-fed) are shown, and the metabolites are grouped by the pattern (HiAlc *Kpn-fed* vs. EtOH-fed) of increased level in different stages.

**Figure S4.**
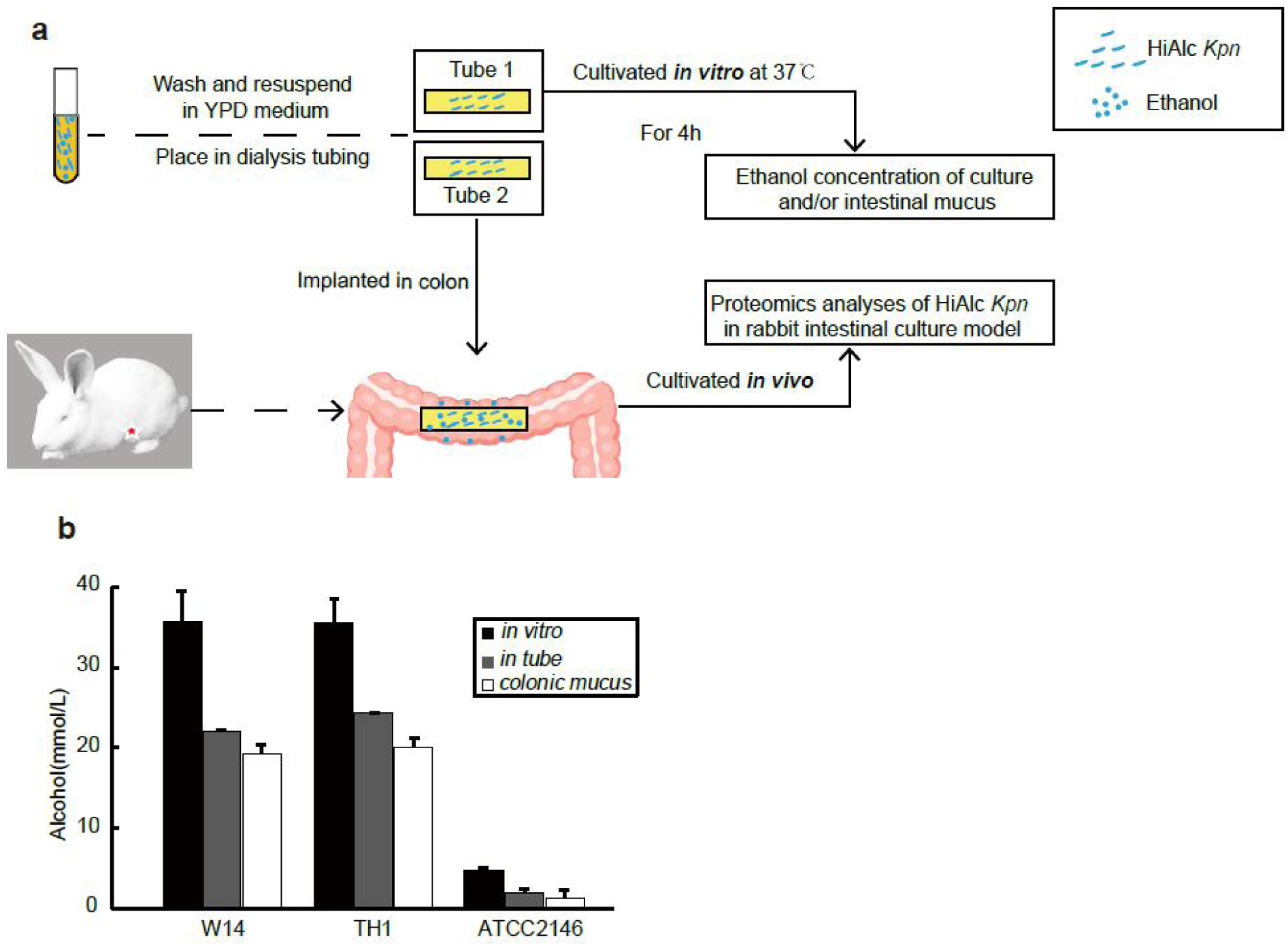
The model for rabbit intestinal culture and proteomic analysis of HiAlc *Kpn* in the in vivo and in vitro conditions. **(a)** Schematic presentation of the rabbit intestinal culture model. **(b)** Alcohol concentration of intestinal mucus after HiAlc *Kpn* W14 and TH1 cultured *in vivo* and *in vitro* for 4h.

**Figure S5.**
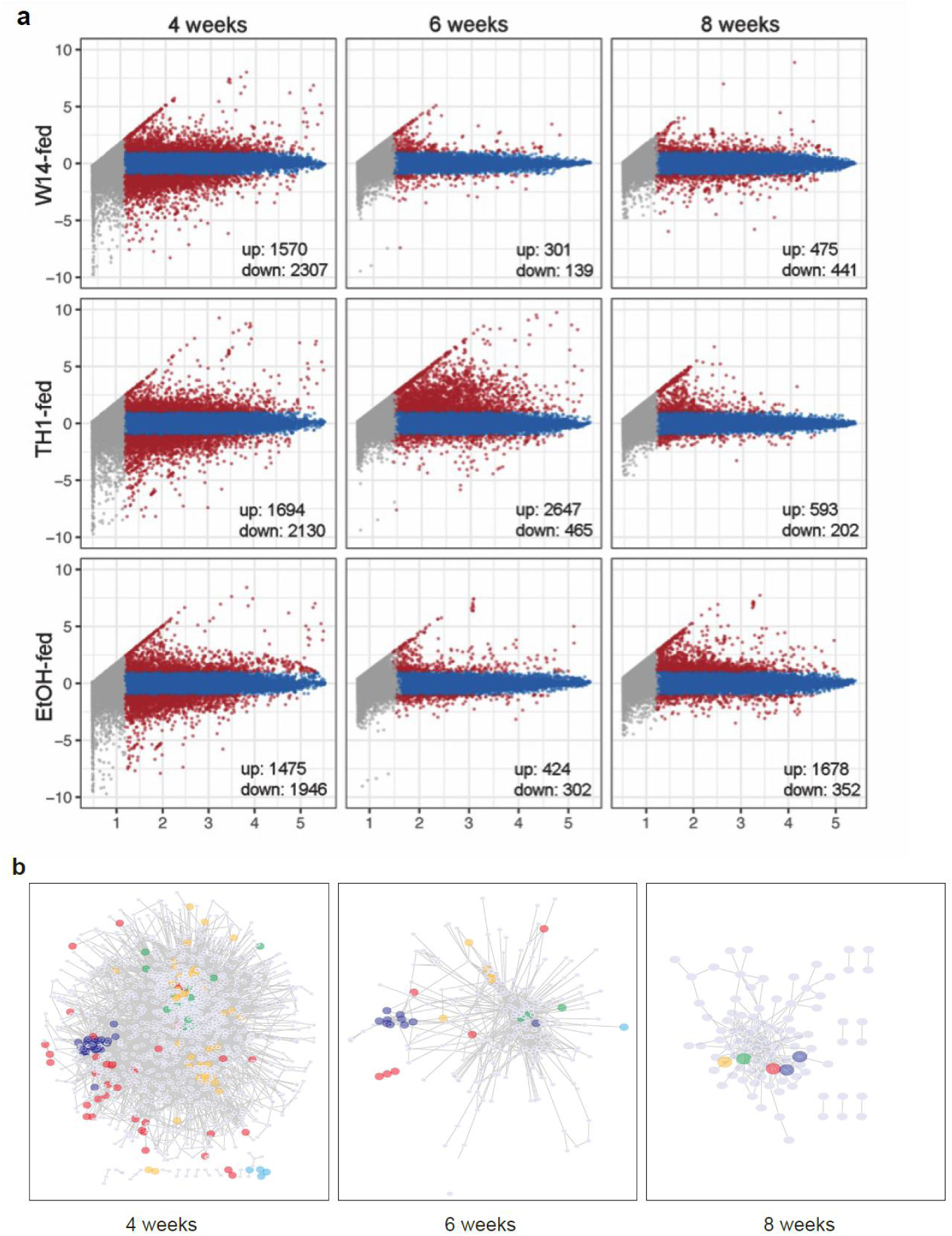
Enrichment of DEGs and Regulatory networks during HiAlc *Kpn* induced FLD development. **(a)**The“Minus-average”(MA) plot to illustrate gene expression profile for pairwise comparison of mice microarray data from HiAlc *Kpn-* and EtOH-fed groups, respectively, against the data from pair-fed group. DEGs (adjusted density ≥ 2 folds to pair-fed group) are shown by red dots, and commonly expressed genes are by blue dots, and non-expressed genes are by grey dots, and commonly expressed genes are by blue dots, and non-expressed genes are by grey dots. The gene number of up and down regulated gene were marked at the right bottom for each panels. **(b)** Regulatory networks of enriched genes after mice fed HiAlc *Kpn* for 4, 6, 8 weeks. We highlight five groups: blue, fatty acid metabolism, including CYP, UGT and HSD gene family; red, alcohol, including ADH, ALDH and SLC gene family; green, immune and inflammatory factors, including IL and SMAD gene family; orange, cancer related factors, including E2F, HIST, RAB and TUBA gene family; and bright blue for Oflr gene family.

**Figure S6.**
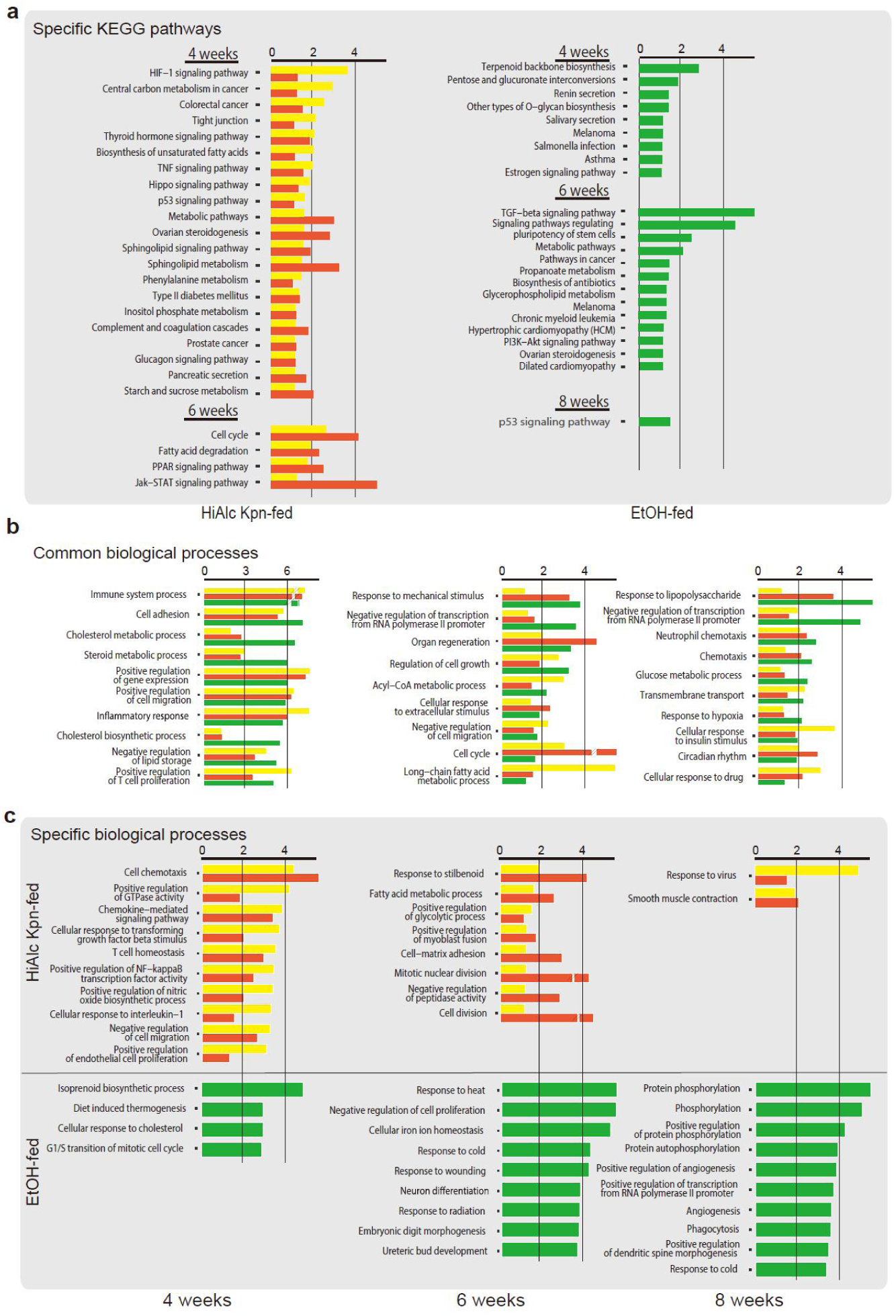
Enriched biological process and KEGG pathway compared to pair-fed groups. **a**, Specifically enriched KEGG pathways in HiAlc *Kpn-* and EtOH-fed mice. **b**, Enriched biological processes in both HiAlc *Kpn-* and EtOH-fed mice. **c**, Specifically enriched biological processes in HiAlc *Kpn-* and EtOH-fed mice.

**Figure S7.**
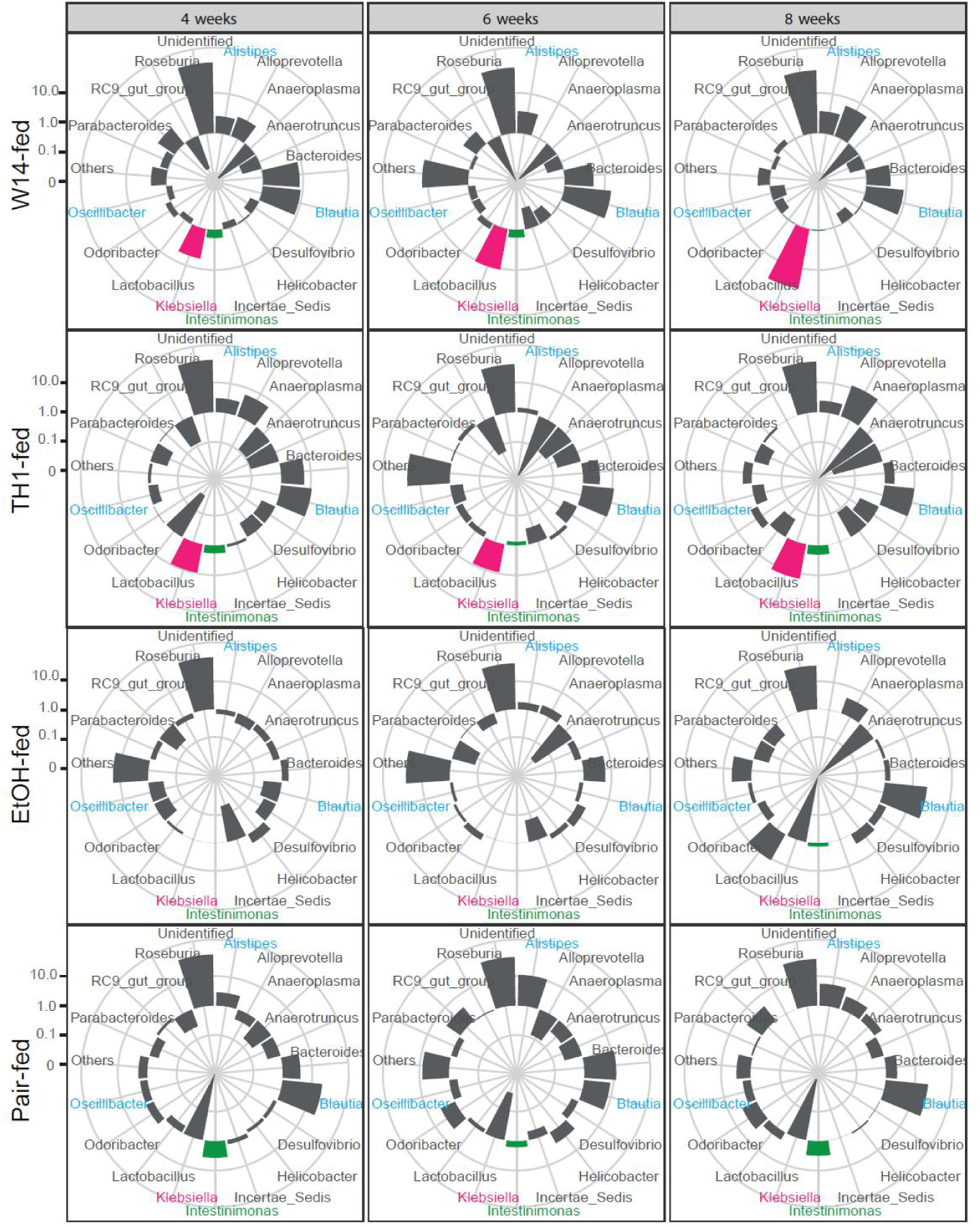
Analyses of intestinal microbiota in FLD mice at 4, 6 or 8 weeks.

